# Easy-to-set-up image analysis characterizes phenotypic diversity in the growth of mushroom-forming fungus *Schizophyllum commune*

**DOI:** 10.1101/2024.06.08.598095

**Authors:** Hiromi Matsumae, Megumi Sudo, Tadashi Imanishi, Tsuyoshi Hosoya

**Affiliations:** Department of Molecular Lifesciences, School of Medicine, Tokai University; Graduate School of Medicine, Tokai University; National Museum Nature and Science (Kahaku), Japan

## Abstract

*Schizophyllum commune*, a common wood-decay mushroom known for its extremely high genetic diversity and as a rare cause of human respiratory diseases, could be a promising model fungus contributing to both biology and medicine. To better understand its phenotypic diversity, we developed an image analysis system that quantifies whole morphological traits of mycelia in Petri dishes. This study evaluated growth of six wild and one clinical isolates of Japanese *S. commune*, subjected to different temperatures and glucose concentrations, including a condition mimicking the human respiratory environment. Our analysis revealed that combinations of two growth indices, area and whiteness, highlighted strain-specific responses, with profiling growth patterns using clustering algorithms. Notably, the clinical isolate exhibited the strongest whiteness under the respiratory-like condition. We also found that the growth rate was strongly determined by glucose concentration, while the effects of temperature on growth varied among the strains, suggesting that while glucose preference is common in this species, responses to temperature differ between strains. Our results suggest that the system possesses sufficient sensitivity to detect growth traits of mycelia. This study provides a key to unravelling unknown traits behind the high polymorphisms in *S. commune*, including the ability to colonize the human respiratory tract.

## Introduction

Although the fungal kingdom harbours phylogenetically six to eight major phyla (Li et al., 2021) and is estimated to host approximately six times the number of species found in terrestrial plants(Kew, 2018), current genomic and molecular biology research has predominantly concentrated on a narrow selection of model organisms in fungi within the Ascomycota phylum. This selection includes yeasts, *Neurospora*, and *Aspergillus*, which exhibit systematic and trait-based biases compared to the diverse model organisms in animal and plant research(McCluskey & Baker, 2017). For example, plant research utilizes a broader range of model organisms, from *Arabidopsis* to rice and poplar trees, tailored to specific study objectives such as breeding, development, ecology, and evolution(Cesarino et al., 2020). Recent advancements in omics and genome editing technologies have accelerated research on a revisit of non-model plants(Cesarino et al., 2020). However, the diversification of fungal studies using these advanced technologies is notably lagging.

Our understanding of the fungal kingdom might be underrepresented, necessitating the integration of various fungal models for both fundamental and applied research into the molecular mechanisms and evolution of fungal traits. *Schizophyllum commune* is a classic model organism in fungal biology(McCluskey & Baker, 2017; MILES, TAKEMARU, & KIMURA, 2006; Raper, Krongelb, & Baxter, 1958) and is known to be distributed throughout the world(Cooke, 1961; Raper et al., 1958; Taylor, Turner, Townsend, Dettman, & Jacobson, 2006). *Schizophyllum commune* is a common wood-decay mushroom, which prefers fresh logs and is also a weak pathogen for living trees(TAKEMOTO, NAKAMURA, IMAMURA, & SHIMANE, 2012). A draft genome of *S. commune* has been sequenced with a size typical for fungi, approximately 38.5-40 Mb(Mohanta & Bae, 2015; Ohm et al., 2010). Population genomics analysis shows that the genetic diversity within the species is extremely high, with a sequence identity of 75-92% between strains from North America, Europe and East Asia(Marian et al., 2024). On average, 14.5 amino acid substitutions per gene were observed within a U.S. population of *S. commune*, which is equivalent to ten times that observed in *Drosophila melanogaster*, making it the most polymorphic species in known eukaryotes(Baranova et al., 2015). The high amino acid level genetic diversity in *S. commune* suggests that there could similarly be high phenotypic diversity to adapt to global environments. Despite the known high genetic diversity within *S. commune*, its diversity of phenotypic traits have been studied under limited conditions such as the development of fruit bodies(Marian et al., 2024), and the link between genes and phenotypes remains poorly understood.

The broad environmental adaptability of *S. commune* might be demonstrated by its pathogenicity towards humans. This species is known to occasionally colonize the human respiratory tract and cause a disease known to be Allergic Bronchopulmonary Mycosis (ABPM)(Amitani et al., 1996; Chowdhary et al., 2013; Oguma et al., 2024, 2018). Among fungi identified as causes of ABPM, Japan has the highest number of cases attributed to *S. commune* in a global clinical survey(Chowdhary et al., 2013). In Japan, *S. commune* is the second most common cause, following only the genus *Aspergillus(Oguma et al., 2024, 2018)*. Although not an infectious agent, *S. commune* can colonize the mucosal surfaces of the bronchi and sinuses for long periods(Amitani et al., 1996). Identification of *S. commune* from clinical samples such as mucus can be achieved through serodiagnosis, culture tests, and DNA testing targeting the ITS region(Asano et al., 2021; Buzina, Lang-Loidolt, Braun, Freudenschuss, & Stammberger, 2001; Chowdhary et al., 2013; Won et al., 2012). For most fungi, optimal growth temperatures are between 25-30℃(Dix & Webster, 1995); however, the internal temperature of the human body is 37℃. The ability to survive in relatively high temperatures may cause fungal diseases in humans(Leach & Cowen, 2013). Much remains unknown about the pathogenesis and treatment of ABPM caused by *S. commune* (Oguma et al., 2024), necessitating verification *in vitro* (Chowdhary et al., 2013). Many strains have been isolated from ABPM patients and are provided as a culture collection of clinical isolates by The Research Center for Medical Mycology at Chiba University and are now available through the National BioResorce Project, Japan. If a method to measure fungal traits *in vitro* can be developed, it could provide insights into the molecular mechanisms of ABPM caused by *S. commune*.

Among the life cycle of *S. commune*, mycelial stages could be technically the easiest to capture its traits. The life cycle of *S. commune* consists of spores, primary mycelium, secondary mycelium, and the reproductive organ, the fruit body (Nieuwenhuis & Aanen, 2018; Palmer & Horton, 2006). As a target for measurement of morphological traits, fruit bodies are not practical for analysis in the current techniques since they have complex three-dimensional shapes and alter their shapes when dried. Once a single spore can be isolated from a fruiting body in the laboratory, it allows for the establishment of genetically clonal mycelial cultures as strains similar to other microbial studies. The development of image analysis methods in microbiological research has been remarkably advanced, with many studies that target counting the number of cells or areas and diameters of colonies and biofilm in microbes(Khalil, Legin, Kurek, Perre, & Taidi, 2021; Ryan et al., 2012; Takeuchi et al., 2014; Zhang et al., 2022). The growth of the mycelia of fungi is more difficult to capture than the growth of unicellular microbes because a mycelum forms complex structure on a solid medium(Khalil et al., 2021). Some experimental systems measure the growth patterns of mycelia proliferating from spores through live imaging systems(Khalil et al., 2021; Ulzurrun, Huang, Chang, Lin, & Hsueh, 2019; Zhang et al., 2022). However, live imaging systems require single-spore isolation for each experiment (i.e., the necessity to induce sexual reproduction and cycle generations), and live imaging involves expensive equipment, making it impractical for routine clinical testing. Therefore, simpler methods are needed to explore strain-specific morphological traits in multicellular fungal cultures.

This study aims to develop an image analysis system to easily detect the whole morphological traits of *Schizophyllum commune*’s mycelia *in vitro* for contributing to biological and medical studies. We prepared four environmental conditions by varying temperature and glucose concentrations, including one condition considered to be closest to the human bronchial surface at 37°C with low glucose. Under these four conditions, mycelia of one clinical isolate and six wild strains from Japan were cultivated on Petri dishes. Images of the Petri dishes were taken on the fist day of transplantation of mycelia and on the fourth day post-transplantation. From these images, two indices of mycelial growth, area and whiteness were measured and growth rates between day 0 and day 4 were calculated. Through statistical analysis and clustering of the growth rates, strain-specific and common growth patterns were captured among the seven Japanese strains of *S. commune*.

## Results

We collected fruit bodies from five geographically distinct locations across Japan and established fungal strains (Fig. 1a-b). These five locations span three of the Köppen climate classifications(Beck et al., 2018) : Akita (FC8125) belongs to the humid continental climate (Df), while Minami-Torishima in the Ogasawara Islands (FC8191-8192) belongs to the savanna climate (Aw), and the remaining three locations belong to the humid subtropical climate (Cfa). Additionally, we included a clinical isolate (IFM 65656) from the Research Center for Medical Mycology at Chiba University, bringing the total to seven strains, which were then cultivated under four conditions and observed on day four (Fig. 1b). The four conditions consisted of two temperature conditions (room temperature and 37°C, which is similar to human body temperature) and two glucose concentration conditions (3.9% PDA and 0.1% PDA) (Experimental Procedures).

**Fig. 1.**
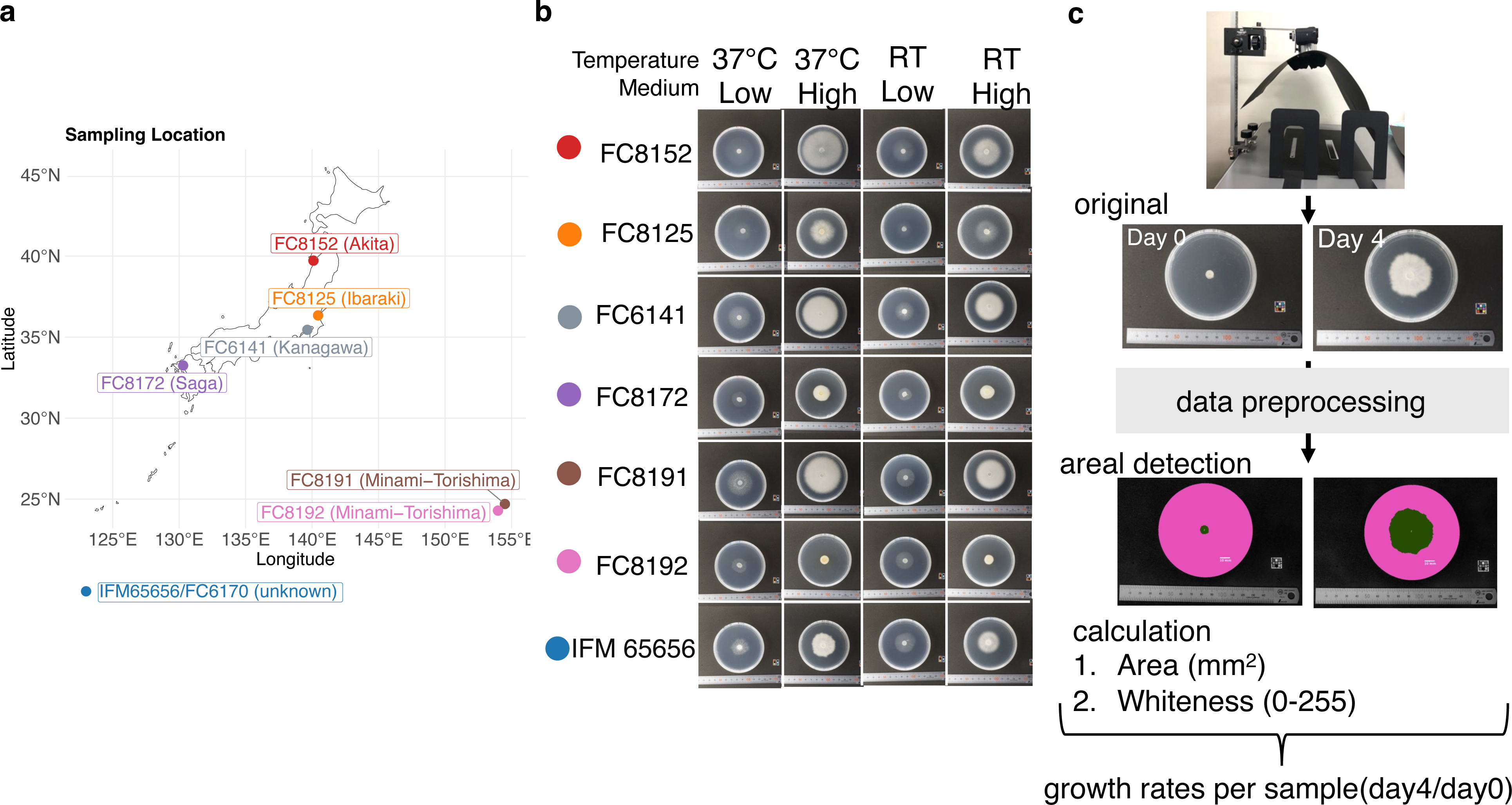
Detecting whole morphological traits of seven strains in *Schizophyllum commune* by image analysis. (a) Geographic origins of the six wild strains. Each color represents a geographic area, except for IFM65656(FC6170) which was isolated from an anonymized patient and, therefore, lacks location. (b) Mycelia cultures observed on Day 4. The horizontal axis lists the strain IDs and locations, and the vertical axis shows combinations of temperature and medium concentration. ‘RT’ indicates Room Temperature. Medium concentration is categorized as Low (0.1% PDA) or High (3.9% PDA). (c) Workflow of image analysis system in this study. Images of mycelia cultured on plastic dishes were captured using a digital camera and processed by ImageJ. The data preprocessing includes color-to-grayscale conversion and background correction. An area of the mycelia (a green section) was separated from the dish and media (a pink section) and measured in square millimeters (mm²). Whiteness is average of grayscale levels ranging from 0 to 255 within the detected area (green). Growth ratios compare areas and whiteness on day 4 to day 0.

In all strains, the mycelia were thin and spread out under low glucose conditions, whereas under high glucose conditions, the mycelia densely spread and became whiter (Fig. 1b). From this observation, whiteness was also suggested as an index of growth, while area and/or diameter have been traditionally used as a growth index in fungal studies (Khalil et al., 2021; Ryan et al., 2012). We observed clear differences between strains in terms of area and whiteness of mycelial growth even on the same glucose concentration (for instance, see 37°C-High in Fig. 1b). Despite both FC8191 and FC8192 being collected from the same 1.51 km² Pacific island, Minami-Torishima, their growth patterns differed remarkably; FC8191 had the largest area among the seven strains, while FC8192 had the smallest (see, for example, 37°C-High and RT-High in Fig. 1b). This suggests that there is no direct relationship between the geographical origin of strains and its growth pattern.

To quantify the observed differences between strains and cultivated conditions, we established an experimental system to compare the growth of mycelia through images (Fig. 1c, Experimental Procedures). Petri dishes were captured from above with a digital camera on the day of inoculation and on day 4. After preprocessing the original images, we obtained an area of the mycelium and average whiteness within that area for each image (Experimental Procedures; Table S1). Growth rates, defined as the ratio of day 4 to day 0, were compared for both area and whiteness across strains and culture conditions (Fig. 2, Table S2). Among the seven strains, FC8172 did not show a statistically significance difference in area across four culture conditions (ANOVA, F(3, 16) = 0.498, p-value = 0.689), whereas the six strains showed significant differences in area under the four conditions(ANOVA, p-value < 0.05)(Fig 2a, Table S3). Additionally, subsequent Tukey’s honestly significant different (HSD) post-hoc comparisons indicated that there were no significant differences in pairwise comparisons among the different conditions in FC8172 (adjusted p-values > 0.05 in all six comparisons, Table S4-5). Of the six strains that showed differences in growth across culture conditions, all but FC8192 tended to have greater areas on high-glucose media (grey shadow in Fig. 2) compared to low-glucose media. The changes in area with temperature were mixed, with some strains growing better at 37°C compared to room temperature. For example, FC8152, collected from Akita, had the largest area at 37°C.

**Fig. 2.**
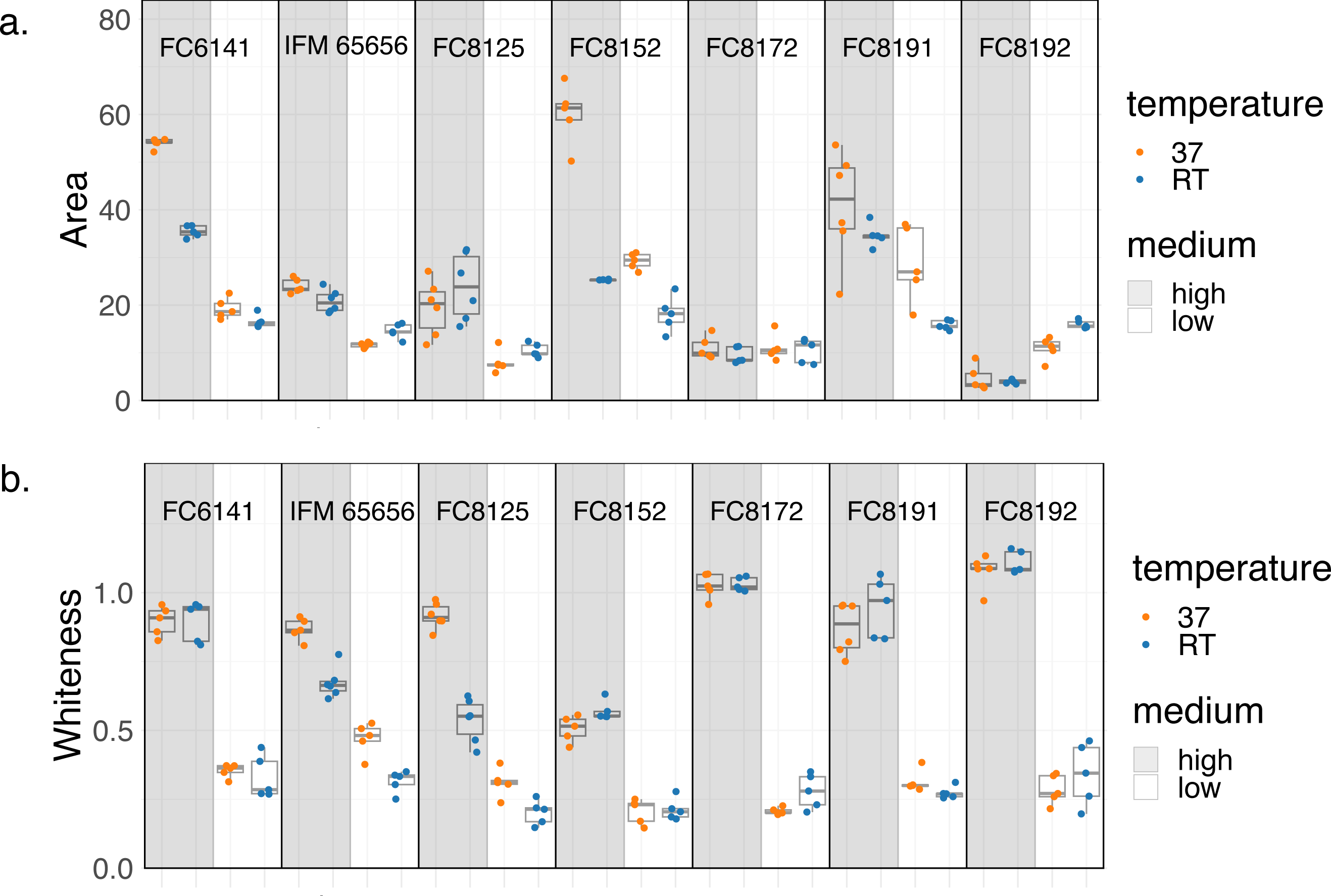
Growth rates of seven strains under four culture conditions. Boxplots show (a) ratios of sizes and (b) whiteness on Day 4 relative to Day 0. Dots are replicates under a specific culture condition. Colors indicate temperature conditions, with orange representing 37°C and blue indicating room temperature (RT). Gray shaded areas denote high-concentration medium (3.9% PDA), while white areas indicate low-concentration medium (0.1% PDA).

Whiteness, on the other hand, generally tended to be lower on day 4 compared to day 0, meaning the mycelium turned greyer over time (Fig. 2b). Unlike area, no strain showed the same growth pattern across all cultivation conditions (ANOVA, p-value < 0.05, shown in Table S2). Notably, FC8172, which showed no significant differences in area across conditions, exhibited clear differences in whiteness (Fig. 2b). Both FC8191 and FC8192, obtained from Minami-Torishima, reflected differences in area in response to changes in cultivation conditions but exhibited similar trends in whiteness. Interestingly, the clinical isolate IFM65656 had the highest whiteness among all strains in the low glucose at 37°C (Fig. 2b). An analysis of the differences between strains across culture conditions using Tukey’s HSD test revealed that IFM65656 and FC8125 exhibited different levels of whiteness in all combinations (adjusted p-values < 0.05; Table S4). For the other five strains, no significant influence of temperature on whiteness was observed within the same medium concentration (Table S4). Taken together, these results yielded different mycelial responses in terms of area and whiteness.

In order to focus on characteristics of mycelia, we examined the relationship between area and whiteness, separated by strain, temperature, and glucose concentration (Fig. 3). Replicates for each strain and culture conditions tended to be in close proximity to each other (Fig. 3a, Fig. S1). The lack of a proportional relationship between area and whiteness suggests that these two indices may reflect different aspects of growth. Area and whiteness were distinctively split based on the glucose level, but temperature did not explain the two indices (Fig. 3b-c). These results suggest that the glucose concentration strongly reflects *S. commune*’s preference for nutrinants, while temperature may reflect traits differences between strains.

**Fig. 3.**
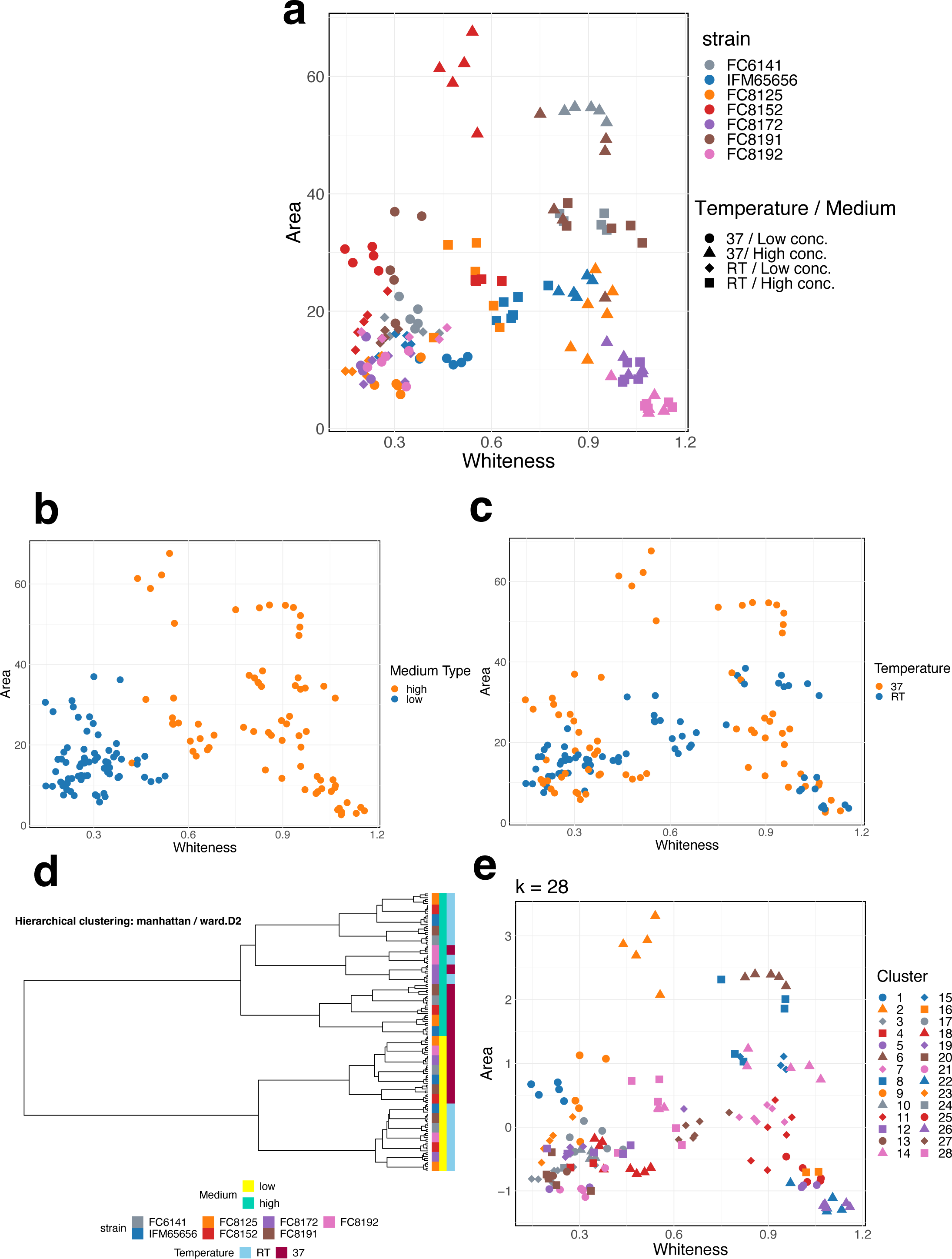
Relationships between areas and whiteness as growth indices in *S. commune.* The Y-axis indicates areas of mycelia, while the X-axis represents mean whiteness within an area. (a) Colors correspond to different strains. Shapes denote combinations of culture conditions, specifically temperature and medium concentration: temperature is either 37°C or room temperature (RT), and medium concentration is categorized as low (0.1% PDA) or high (3.9% PDA). (b)The plot from (a) is differentiated by medium concentration; orange for high concentration and blue for low concentration. (c) The plot from (a) is distinguished based on culture temperature; orange for 37°C, and blue for room temperature. (f) Hierarchical clustering with information on area, whiteness, strain, temperature, and glucose level was provided. A dendrogram was constructed by the Manhattan distance and Ward’s method. The color bars on the right side of the dendrogram categorize the samples by strain, medium concentration, and temperature. (g) K-means clustering with 28 clusters for the same data from (f). The color and shape of each point correspond to its unique cluster number.

Next, we classified the area and whiteness by combining information on strain, temperature, and glucose concentration using two clustering methods (Fig. 3d-e, Experimental Procedures). The scatter plot in Figure 3a should be able to classify the data into 28 clusters, representing the combinations of 7 strains and 4 culture conditions. In hierarchical clustering, all biological replicates for each combination of strain and condition formed a single cluster (Fig. 3d). The dendrogram initially split perfectly by glucose level, reflecting the observed data (Fig. 3b). Although the dendrogram tended to split at the second tier by temperature, the strains FC8172 and FC8192 (purple and pink in Fig. 3d), which showed little change in area at high glucose (Fig. 2a), could not be differentiated based on their responses to temperature. The accuracy of classification was reduced when information on strain ID was removed from the dataset (Fig. S2), suggesting that including information about biological replicates leads to improved clustering. Applying the K-means method with K=28 to the same data, replicates tended to be labelled the same colour (Fig. 3e). For example, it successfully classified FC6141 and FC8152 under 37℃ and high glucose (grey and red triangles in Fig 3a) into clusters 2 and 6, respectively. The clustering was also reconstructed under room temperature and low glucose; FC8152 (red diamond in Fig. 3a) was mapped into cluster 23 (orange diamond in Fig. 3e). When K-means clustering with K=2 was applied, it yielded a different classification result compared to hierarchical clustering, which initially split the data based on glucose level (Figure S3).

## Discussion

In this study, we developed the image analyzing system to capture the characteristics of mycelial growth using seven strains of Japanese *Schizophyllum commune*, aiming to utilize this species as a fungal model in molecular biology and medical research. Although the shape of the fruit bodies is three-dimensionally complex, capturing the overall image of the mycelia in two dimensions allowed us to highlight the differences in growth among the strains. This method, which uses the common digital camera and culture media, is cost-effective and straightforward as it does not require specialized equipment or experimental procedures (Fig. 1c). We believe this method is suitable for screening the morphological traits of *S. commune* when incorporating genomic science and molecular biological techniques.

While our system captures macroscopic features of the mycelia, enhancing its quantitativeness requires capturing microscopic-level characteristics. For instance, mycelia that are translucent and difficult to automatically detect in terms of area may have low cell density, which can be verified by counting cell numbers under a microscope. In fungi, some methods that quantify proliferation from a single spore have already been reported (Khalil et al., 2021; Ulzurrun et al., 2019; Zhang et al., 2022). However, we could not use these previous methods since we transplanted sections of mycelium that had proliferated from a single spore and thus took replicates from clones. Cell counting in mycelium is challenging because, unlike bacteria, fungal mycelium grows in a three-dimensional manner, like a ball of yarn, making it difficult to count the number of cells from two-dimensional images. Future development of microscopic methods that enable the counting of both initially transplanted and subsequently grown mycelial cells would capture exact growth rates at the cellular level.

Our study shows that whiteness could be the new index to measure the growth of mycelium as well as conventional indices like area and diameters in fungal studies(Khalil et al., 2021; Ryan et al., 2012). The proportional relationship was not observed between area and whiteness and combinations of area and whiteness identified strain-specific growth, suggesting that area and whiteness may serve as distinct indices of *S. commune* growth. By combining these two measures with cultivation conditions such as temperature and glucose concentration, it may be possible to automatically identify strain-specific profiles (Fig. 3d-e). Interestingly, while the area of FC8172 remained constant across different cultivation conditions, its whiteness varied with glucose level (Fig. 2), suggesting that whiteness may reflect a strain-specific trait for FC8172. The biological replicates indicated the presence of strains like IFM65656, which showed little variation in area and whiteness, and different types of strains like FC8191, which exhibited a large variety (Fig. S1). Whether these observed variations among strains are due to biological or technical reasons will need to be investigated further. Furthermore, the opposite growth patterns were displayed by FC8191 and FC8192, obtained from Minami-Torishima (Fig. 2a), suggesting that morphological growth patterns may not reflect phylogeographic distinctions within the Japanese archipelago.

The common response among the strains was observed in relation to the glucose concentration in the mediua, which affected both the area and whiteness (Fig. 3b). This observation may reflect the ecological preference of *S. commune* in natural environments, which is known to colonize sugar-rich early decaying wood or to invade living tree bark (Marian et al., 2024; TAKEMOTO et al., 2012). This trait is supported at the genetic level, as the species have a set of specific enzymes for decomposing the bark of living trees, —a defense system of the trees—as well as a set of enzymes for degrading dead wood(Almási et al., 2019). Thus, genes related to the response to glucose might be evolutionary more conservative than those related to other responses within the species.

In contrast to glucose concentration, the temperature-induced changes in growth response to temperature varied among strains. It might be a key to understanding whether *S. commune* can colonize within the human body. Under low glucose at 37°C, IFM65656, isolated from an ABPM patient, exhibited the strongest whiteness among the seven strains tested. This result may indicate that *S.commune* populations harbour high genetic diversity *in natura*, with strains capable of responding to various environments, including some that may easily colonize the human body. However, this analysis compared only one clinical isolate, IFM65656, against six wild strains, which is a limited sample size. Future studies should increase the number of clinical isolates compared to provide a comprehensive analysis of the phenotypic diversity of *S. commune*. While the draft genome of a strain *S. commune* isolated from North America has been sequenced(Ohm et al., 2010) and the global genetic diversity has been studied(Baranova et al., 2015; Marian et al., 2024; Taylor et al., 2006), the genetic diversity of Japanese populations, including amino acid polymorphisms, is still not well understood. In the future, by analyzing the relationship between genetic diversity and phenotypic diversity, it may be possible to contribute to elucidating the pathogenesis and treatment methods of ABPM caused by *S. commune*, which occurs most frequently in Japan (Asano et al., 2021; Chowdhary et al., 2013; Oguma et al., 2024, 2018).

Our analysis would also contribute to understanding the relationships between genetic and phenotypic diversity in nature. The recent report illuminates how intraspecific variation in animals, plants, and also fungus affects ecosystems(Roches et al., 2018). *Schizophyllum commune* is a common wood decomposer across continents, and it is well-represented in the world’s largest public biodiversity database, the Global Biodiversity Information Facility (GBIF)(GBIF, 2024), with over 70,000 observation records as of April 17, 2024. The role of *S. commune*’s high genetic diversity in its phenotypic diversity in the wild, such as altering the amount and speed of wood decomposition, or its impact on forest biomass, remains unclear. Investigating the intraspecific diversity of *S. commune* could be intriguing not only from a medical standpoint but also in terms of understanding the evolution of its diverse genome and its impact on ecosystems.

## Experimental procedures

### Fungal Culture

We used seven strains of *S. commune*, including a clinical strain isolated from an ABPM patient stored in the Research Center for Medical Mycology, Chiba University (ID: IFM 65656, managed in this study as FC6170) and six wild isolates from different geographic origins archived in the fungal collection of the National Museum of Nature and Science (Kahaku), Japan (Fig. 1). The wild strains were cultured as single-spore isolates from fruiting bodies, either as primary or secondary mycelia. *Schizophyllum commune* has a life cycle that includes both primary and secondary mycelial states, distinguishable by the presence or absence of a characteristic cellular structure called clamp connections. Among the seven strains used in this study, FC8125 was identified as a secondary mycelium due to the presence of clamp connections upon microscopic observation, while the remaining six strains were considered primary mycelia due to the absence of clamp connections. All the wild isolates can be provided upon request.

Cultivated conditions were designed to create both human-like and non-human-like environments by varying the concentration of the medium and the incubation temperature. Potato dextrose agar (PDA, Nissui Pharmaceutical Co.) was prepared at high (3.9%) and low (0.1%) concentrations, and 10 ml of each was dispensed into 90 mm sterile Petri dishes. *Schizophyllum commune* favours fresh wood in its natural habitat(Almási et al., 2019; TAKEMOTO et al., 2012), while the respiratory tract (on the bronchial surface) may lack sufficient major nutrients for fungi, like glucose. We employed 0.1% of PDA as a “low” glucose medium to mimic fasting blood glucose levels of 0.1%. A fragment of mycelia with the medium was extracted by An 8 mm cork borer, and then it was inoculated at the centre of a new dish for each experiment. As the human interior (such as the sinuses and bronchi) is a dark environment, all cultures were grown in dark conditions. Incubation temperatures were set at room temperature (to simulate a natural environment) and 37°C (to simulate human body temperature). Therefore, the combination of a low-concentration glucose medium and 37°C incubation was deemed the closest approximation to a human body-like environment in four possible combinations of medium and temperature. For image analysis, 3-5 replicates per condition were prepared, utilising mycelia from both day 0 and day 4.

### Image Analysis

A digital camera was mounted directly above a Petri dish at a distance of about 44 cm from the lens to the desk, and a black paper was placed over the top to prevent reflections from the dish lid (Fig. 2). To maintain safety, photos of the whole dishes were taken with the lid on, under the black paper. All the original images are available from 10.5281/zenodo.11180775.

Image J 1.53k (Schneider et al., Nature Methods, 2012) was used to preprocess the images to control for lighting effects. First, background correction was performed using the rolling ball algorithm (ball radius = 1500 pixels) to subtract the background. Then, the background-subtracted images were converted to 8-bit grayscale, reducing them from colour images to 256-level grayscale images, where values closer to 0 indicate dark and 255 indicate white. The camera and subject positions were fixed, ensuring that the scale did not change between images, with the scale set at approximately 26.5 pixels per millimetre.

Next, to detect growth indices specific to each strain, namely area and whiteness, the mycelial outline was detected in one of the following ways. The area of the outline (in mm²) was computed by ImageJ. The whiteness level was defined by the mean grey value (0-255) within the outline. Area and whiteness for all samples were shown in Table S1.

1. In the cases of mycelia that grew entirely white, the region was detected automatically (particle size 50 pixels, circularity 0-1) (Fig. 2).
2. For mycelia that were generally faint and showed little change between day 0 and day 4, the outline was obtained by subtracting the day 0 image from the day 4 image by Image Calculator. The subtracted image was then converted to 8-bit grayscale, scaled, and thresholded with the mean dark algorithm for particle analysis. Dishes were then encircled with an ellipse, and particle analysis was conducted to detect outlines.
3. For the remaining images where neither of these methods worked, freehand tools were used to outline the mycelium manually.

### Statistical Analysis

Data visualisation, statistical analyses, and clustering of area and whiteness were carried out in R 4.3.1(Team, 2023) using the ggplot2(Wickham, 2016), dplyr(Wickham, François, Henry, Müller, & Vaughan, 2023), tidyr(Wickham, Vaughan, & Girlich, 2023), and broom(Robinson, Hayes, & Couch, 2023) packages. Some parts of the source codes were written by ChatGPT v4(AI, 2023). The in-house R scripts can be accessed on GitHub, https://github.com/mhiromi/scom_mycelial_growth_2024.

For map rendering, the rnaturalearth(Massicotte & South, 2023) and ggrepel(Slowikowski, 2024) packages were used. Growth rates were defined as the ratio of day 4 to day 0 in terms of area and whiteness (Table S2), and these rates were utilised in subsequent statistical analyses.

The differences in area and whiteness across four culture conditions (temperature and medium) for each strain were tested using ANOVA and Tukey’s honestly significant difference (HSD) test from the car(Fox & Weisberg, 2019) and agricolae(Mendiburu, 2023) packages ( Table S3-4). Given the clear differences in area and whiteness based on the medium (Fig 3b), F-statistics and T-statistics were calculated. Mean values and standard deviations between replicates under the same culture conditions for each strain were computed (Fig. S1).

We performed clustering of the data, including the two indices and cultivated information. The data contained both numerical (area and whiteness) and categorical data (temperature, medium, and strain ID), and thus preprocessing was necessary before clustering. Categorical data were converted into binary information using one-hot vector encoding with the fastDummies(Kaplan, 2023) package in R (Table S6). Area and whiteness were then scaled using the `scale` function to address differing scales. Hierarchical clustering was performed using the `hclust` function with the Manhattan distance and Ward.D2 method (Fig. 3, Fig. S2). The obtained dendrograms were rendered using the ape(Paradis & Schliep, 2019) and ggtree(Yu, 2023) packages. K-means clustering was performed using the `kmeans` function (Fig. 3, Fig. S3). K=28 was set as the optimal cluster number to combine all strain, medium, and temperature conditions.

## Acknowledgements

The authors thank Kentaro Hosaka (Kahaku) for providing fruit bodies from Minami-Torishima island. The authors thank Katsuhiko Kamei (Chiba Univ), Yoshiki Shiraishi (Tokai Univ), Takeru Nakazato (DBCLS), and Mika Sakamoto (NIG) for the discussion. Clinical isolate IFM 65656 was provided by Medical Mycology Research Center, Chiba University, with support in part by National BioResource Project (NBRP), AMED, Japan (https://nbrp.jp/). A part of this work was performed using the facilities of the Medical Science College Office, Tokai University.

This work was partly supported by MEXT KAKENHI Numbers 20K16254, 21H04358 to H.M, ROIS-DS-JOINT-GRANT (029RP2019) to H.M, Joint Usage/Research Program of Medical Mycology Research Center, Chiba University (19-11) to H.M, 2019-2020 Tokai University School of Medicine Research Aid to H.M. Author contributions: HM and TI designed the research. TH isolated strains from fruit bodies. TH and MS cultured *S. commune*. MS captured images. HM and MS processed image data and performed statistical analyses. All authors reviewed the final manuscript.

## Notes

https://zenodo.org/records/11180775

https://github.com/mhiromi/scom_mycelial_growth_2024

### Competing Interest Statement

The authors have declared no competing interest.

## References

AI, O. (2023). ChatGPT v4. Retrieved 2024, from ChatGPT v4 website: https://www.openai.com/chatgpt

Almási, É., Sahu, N., Krizsán, K., Bálint, B., Kovács, G. M., Kiss, B., … Nagy, L. G. (2019). Comparative genomics reveals unique wood-decay strategies and fruiting body development in the Schizophyllaceae. New Phytologist, 224(2), 902–915. doi: 10.1111/nph.16032

Amitani, R., Nishimura, K., Niimi, A., Kobayashi, H., Nawada, R., Murayama, T., … Kuze, F. (1996). Bronchial Mucoid Impaction Due to the Monokaryotic Mycelium of Schizophyllum commune. Clinical Infectious Diseases, 22(1), 146–148. doi: 10.1093/clinids/22.1.146

Asano, K., Hebisawa, A., Ishiguro, T., Takayanagi, N., Nakamura, Y., Suzuki, J., … Program, J. A. R. (2021). New clinical diagnostic criteria for allergic bronchopulmonary aspergillosis/mycosis and its validation. Journal of Allergy and Clinical Immunology, 147(4), 1261–1268.e5. doi: 10.1016/j.jaci.2020.08.029

Baranova, M. A., Logacheva, M. D., Penin, A. A., Seplyarskiy, V. B., Safonova, Y. Y., Naumenko, S. A., … Kondrashov, A. S. (2015). Extraordinary Genetic Diversity in a Wood Decay Mushroom. Molecular Biology and Evolution, 32(10), 2775–2783. doi: 10.1093/molbev/msv153

Beck, H. E., Zimmermann, N. E., McVicar, T. R., Vergopolan, N., Berg, A., & Wood, E. F. (2018). Present and future Köppen-Geiger climate classification maps at 1-km resolution. Scientific Data, 5(1), 180214. doi: 10.1038/sdata.2018.214

Buzina, W., Lang-Loidolt, D., Braun, H., Freudenschuss, K., & Stammberger, H. (2001). Development of Molecular Methods for Identification of Schizophyllum commune from Clinical Samples. Journal of Clinical Microbiology, 39(7), 2391–2396. doi: 10.1128/jcm.39.7.2391-2396.2001

Cesarino, I., Ioio, R. D., Kirschner, G. K., Ogden, M. S., Picard, K. L., Rast-Somssich, M. I., & Somssich, M. (2020). Plant *Science*’s Next Top Models. Annals of Botany, 126(1), 1–23. doi: 10.1093/aob/mcaa063

Chowdhary, A., Randhawa, H. S., Gaur, S. N., Agarwal, K., Kathuria, S., Roy, P., … Meis, J. F. (2013). Schizophyllum commune as an emerging fungal pathogen: a review and report of two cases. Mycoses, 56(1), 1–10. doi: 10.1111/j.1439-0507.2012.02190.x

Cooke, W. B. (1961). The Genus Schizophyllum. Mycologia, 53(6), 575. doi: 10.2307/3756459

Dix, N. J., & Webster, J. (1995). Fungal Ecology. 322–340. doi: 10.1007/978-94-011-0693-1_12

Fox, J., & Weisberg, S. (2019). An R Companion to Applied Regression. Retrieved from https://socialsciences.mcmaster.ca/jfox/Books/Companion/

GBIF. (2024). https://www.gbif.org. Retrieved from https://www.gbif.org

Kaplan, J. (2023). fastDummies: Fast Creation of Dummy (Binary) Columns and Rows from Categorical Variables. Retrieved from https://CRAN.R-project.org/package=fastDummies

Kew, R. B. G. (2018). State of the World’s Fungi 2018 ([“Willis & Katherine”], Eds.). Royal Botanic Gardens, Kew.

Khalil, H., Legin, E., Kurek, B., Perre, P., & Taidi, B. (2021). Morphological growth pattern of Phanerochaete chrysosporium cultivated on different Miscanthus x giganteus biomass fractions. BMC Microbiology, 21(1), 318. doi: 10.1186/s12866-021-02350-8

Leach, M. D., & Cowen, L. E. (2013). Surviving the Heat of the Moment: A Fungal Pathogens Perspective. PLoS Pathogens, 9(3), e1003163. doi: 10.1371/journal.ppat.1003163

Li, Y., Steenwyk, J. L., Chang, Y., Wang, Y., James, T. Y., Stajich, J. E., … Rokas, A. (2021). A genome-scale phylogeny of the kingdom Fungi. Current Biology, 31(8), 1653–1665.e5. doi: 10.1016/j.cub.2021.01.074

Marian, I. M., Valdes, I. D., Hayes, R. D., LaButti, K., Duffy, K., Chovatia, M., … Ohm, R. A. (2024). High phenotypic and genotypic plasticity among strains of the mushroom-forming fungus Schizophyllum commune. BioRxiv, 2024.02.21.581338.doi: 10.1101/2024.02.21.581338

Massicotte, P., & South, A. (2023). rnaturalearth: World Map Data from Natural Earth. Retrieved from https://CRAN.R-project.org/package=rnaturalearth

McCluskey, K., & Baker, S. E. (2017). Diverse data supports the transition of filamentous fungal model organisms into the post-genomics era. Mycology, 8(2), 67–83. doi: 10.1080/21501203.2017.1281849

Mendiburu, F. de. (2023). agricolae: Statistical Procedures for Agricultural Research. Retrieved from https://CRAN.R-project.org/package=agricolae

MILES, P. G., TAKEMARU, T., & KIMURA, K. (2006). Incompatibility Factors in the Natural Population of Schizophyllum commune. Shokubutsugaku Zasshi, 79(940– 941), 693. doi: 10.15281/jplantres1887.79.693

Mohanta, T. K., & Bae, H. (2015). The diversity of fungal genome. Biological Procedures Online, 17(1), 8. doi: 10.1186/s12575-015-0020-z

Nieuwenhuis, B. P. S., & Aanen, D. K. (2018). Nuclear arms races: Experimental evolution for mating success in the mushroom-forming fungus Schizophyllum commune. PLoS ONE, 13(12), e0209671. doi: 10.1371/journal.pone.0209671

Oguma, T., Ishiguro, T., Kamei, K., Tanaka, J., Suzuki, J., Hebisawa, A., … Program, J. A. R. (2024). Clinical characteristics of allergic bronchopulmonary mycosis caused by Schizophyllum commune. Clinical and Translational Allergy, 14(1), e12327. doi: 10.1002/clt2.12327

Oguma, T., Taniguchi, M., Shimoda, T., Kamei, K., Matsuse, H., Hebisawa, A., … Asano, K. (2018). Allergic bronchopulmonary aspergillosis in Japan: A nationwide survey. Allergology International, 67(1), 79–84. doi: 10.1016/j.alit.2017.04.011

Ohm, R. A., Jong, J. F. de, Lugones, L. G., Aerts, A., Kothe, E., Stajich, J. E., … Wösten, H. A. B. (2010). Genome sequence of the model mushroom Schizophyllum commune. Nature Biotechnology, 28(9), 957–963. doi: 10.1038/nbt.1643

Palmer, G. E., & Horton, J. S. (2006). Mushrooms by magic: making connections between signal transduction and fruiting body development in the basidiomycete fungus Schizophyllum commune. FEMS Microbiology Letters, 262(1), 1–8. doi: 10.1111/j.1574-6968.2006.00341.x

Paradis, E., & Schliep, K. (2019). ape 5.0: an environment for modern phylogenetics and evolutionary analyses in R. Bioinformatics, 35, 526–528. doi: 10.1093/bioinformatics/bty633

Raper, J. R., Krongelb, G. S., & Baxter, M. G. (1958). The Number and Distribution of Incompatibility Factors in Schizophyllum. The American Naturalist, 92(865), 221–232. doi: 10.1086/282030

Robinson, D., Hayes, A., & Couch, S. (2023). broom: Convert Statistical Objects into Tidy Tibbles. Retrieved from https://CRAN.R-project.org/package=broom

Roches, S. D., Post, D. M., Turley, N. E., Bailey, J. K., Hendry, A. P., Kinnison, M. T., … Palkovacs, E. P. (2018). The ecological importance of intraspecific variation. Nature Ecology & Evolution, 2(1), 57–64. doi: 10.1038/s41559-017-0402-5

Ryan, O., Shapiro, R. S., Kurat, C. F., Mayhew, D., Baryshnikova, A., Chin, B., … Boone, C. (2012). Global Gene Deletion Analysis Exploring Yeast Filamentous Growth. Science, 337(6100), 1353–1356. doi: 10.1126/science.1224339

Slowikowski, K. (2024). ggrepel: Automatically Position Non-Overlapping Text Labels with “ggplot2.” Retrieved from https://CRAN.R-project.org/package=ggrepel

TAKEMOTO, S., NAKAMURA, H., IMAMURA, Y., & SHIMANE, T. (2012). Schizophyllum commune as a Ubiquitous Plant Parasite. Japan Agricultural Research Quarterly: JARQ, 44(4), 357. doi: 10.6090/jarq.44.357

Takeuchi, R., Tamura, T., Nakayashiki, T., Tanaka, Y., Muto, A., Wanner, B. L., & Mori, H. (2014). Colony-live — a high-throughput method for measuring microbial colony growth kinetics— reveals diverse growth effects of gene knockouts in Escherichia coli. BMC Microbiology, 14(1), 171. doi: 10.1186/1471-2180-14-171

Taylor, J. W., Turner, E., Townsend, J. P., Dettman, J. R., & Jacobson, D. (2006). Eukaryotic microbes, species recognition and the geographic limits of species: examples from the kingdom Fungi. Philosophical Transactions of the Royal Society B: Biological Sciences, 361(1475), 1947–1963. doi: 10.1098/rstb.2006.1923

Team, R. C. (2023). R: A Language and Environment for Statistical Computing. Retrieved from https://www.R-project.org/

Ulzurrun, G. V.-D. de, Huang, T.-Y., Chang, C.-W., Lin, H.-C., & Hsueh, Y.-P. (2019). Fungal feature tracker (FFT): A tool for quantitatively characterizing the morphology and growth of filamentous fungi. PLoS Computational Biology, 15(10), e1007428. doi: 10.1371/journal.pcbi.1007428

Wickham, H. (2016). ggplot2: Elegant Graphics for Data Analysis. Retrieved from https://ggplot2.tidyverse.org

Wickham, H., François, R., Henry, L., Müller, K., & Vaughan, D. (2023). dplyr: A Grammar of Data Manipulation. Retrieved from https://CRAN.R-project.org/package=dplyr

Wickham, H., Vaughan, D., & Girlich, M. (2023). tidyr: Tidy Messy Data. Retrieved from https://CRAN.R-project.org/package=tidyr

Won, E. J., Shin, J. H., Lim, S. C., Shin, M. G., Suh, S. P., & Ryang, D. W. (2012). Molecular Identification of Schizophyllum commune as a Cause of Allergic Fungal Sinusitis. Annals of Laboratory Medicine, 32(5), 375–379. doi: 10.3343/alm.2012.32.5.375

Yu, G. (2023). Data Integration, Manipulation and Visualization of Phylogenetic Trees. doi: 10.1201/9781003279242

Zhang, J., Li, C., Rahaman, M. M., Yao, Y., Ma, P., Zhang, J., … Grzegorzek, M. (2022). A comprehensive review of image analysis methods for microorganism counting: from classical image processing to deep learning approaches. Artificial Intelligence Review, 55(4), 2875–2944. doi: 10.1007/s10462-021-10082-4

