## Supplementary Figure S1-3 for "Easy-to-set-up image analysis characterizes phenotypic diversity in the growth of mushroom-forming fungus *Schizophyllum commune*"

**Fig. S1. Relationships between average area and average whiteness under different culture conditions.** Y-axis indicates an averaged area of mycelia, while the X-axis represents mean whiteness within the area. Colors correspond to different strains. Shapes denote combinations of culture conditions, specifically temperature and medium concentration: temperature is either 37° C or room temperature (RT), and medium concentration is categorized as low (0.1% PDA) or high (3.9% PDA). Error bars at each point indicate standard deviations for each condition.

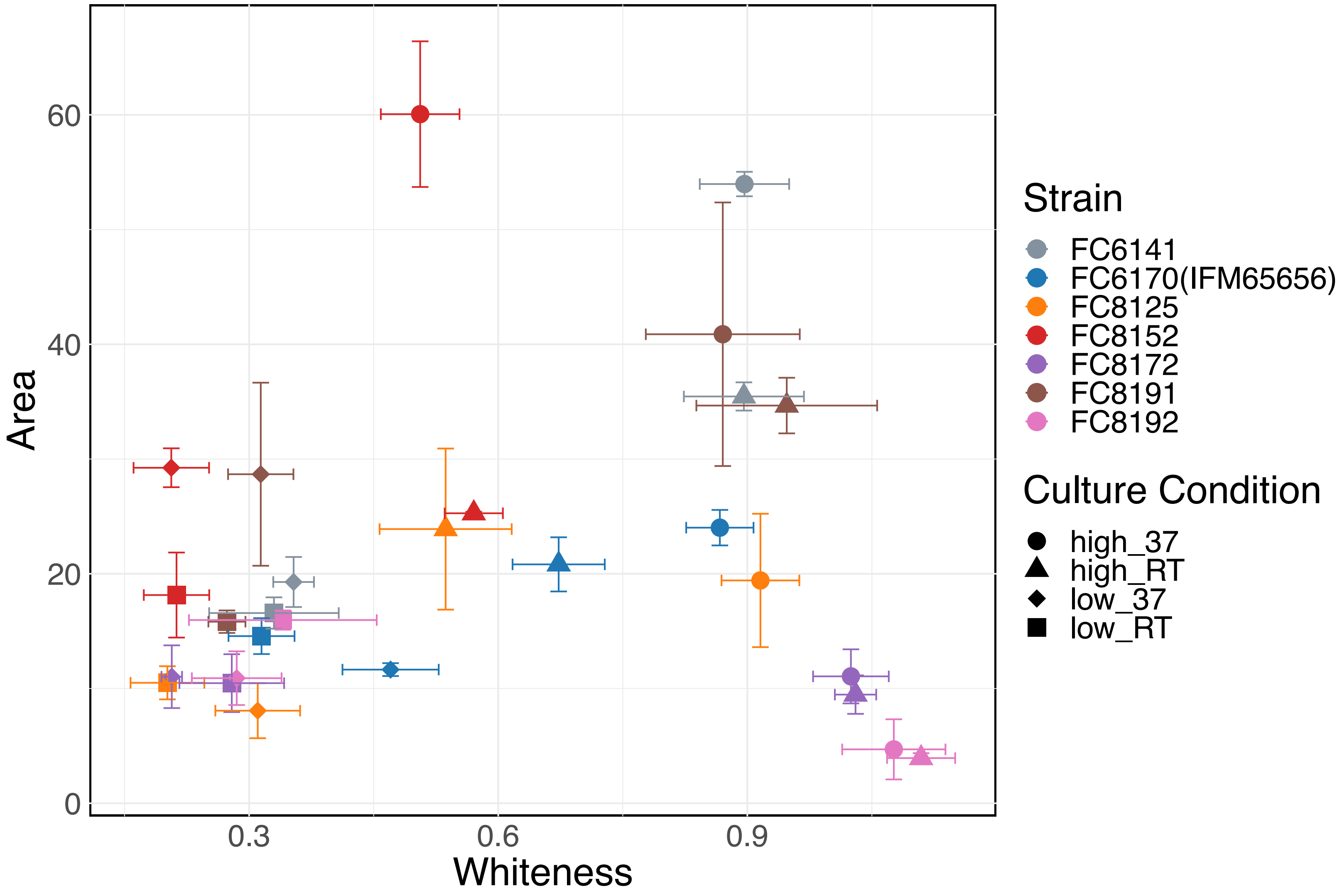

**Fig. S2. Hierarchical clustering without strain information.** This figure displays s hierarchical clustering dendrogram when strain information are absent. Each node in the dendrogram represents a sample. Color bars located to the right of the dendrograms categorize the samples according to strain, medium, and temperature**.**

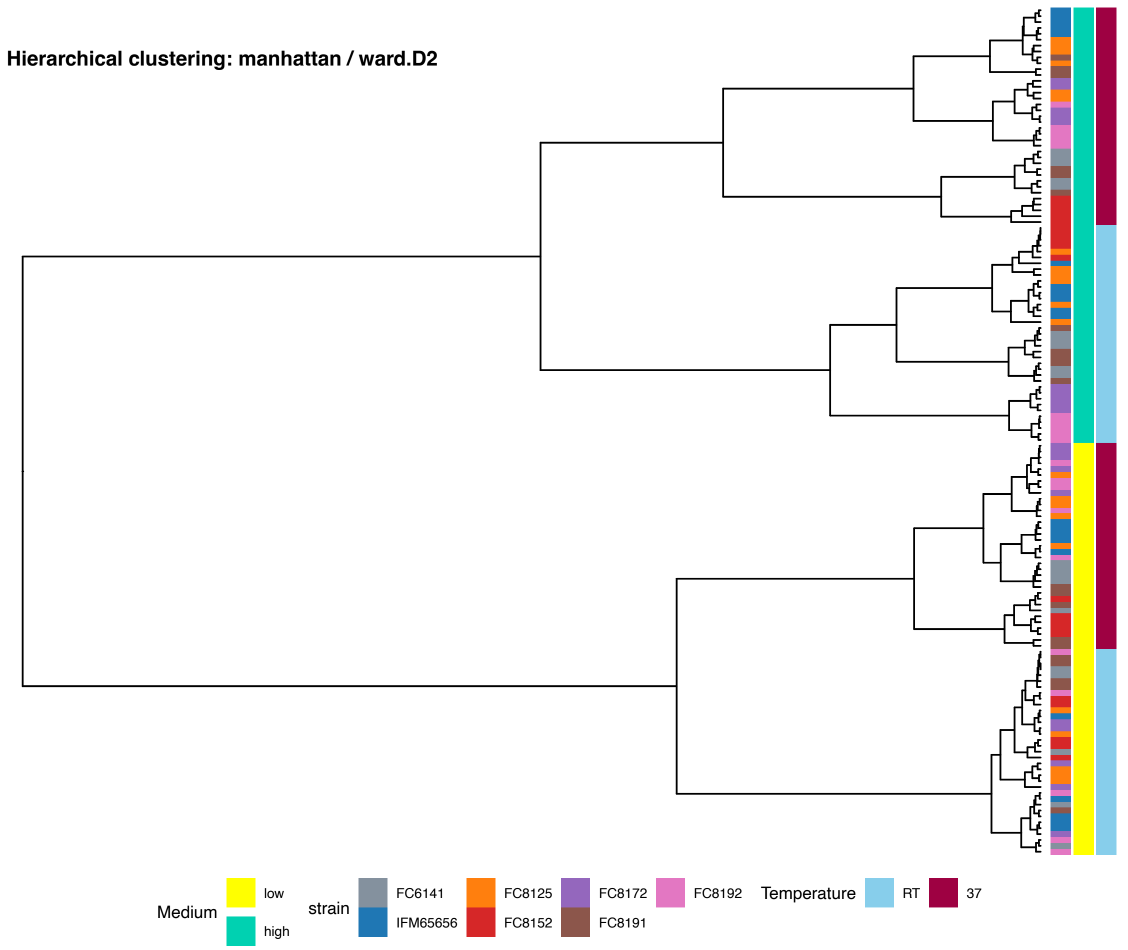

**Fig. S3. K-means clustering of growth information from k=2.** K-means clustering with 2 clusters for the same data from (Fig 3f). The color and shape of each point correspond to its unique cluster number. Combinations color and shape indicate 28 cluster numbers.

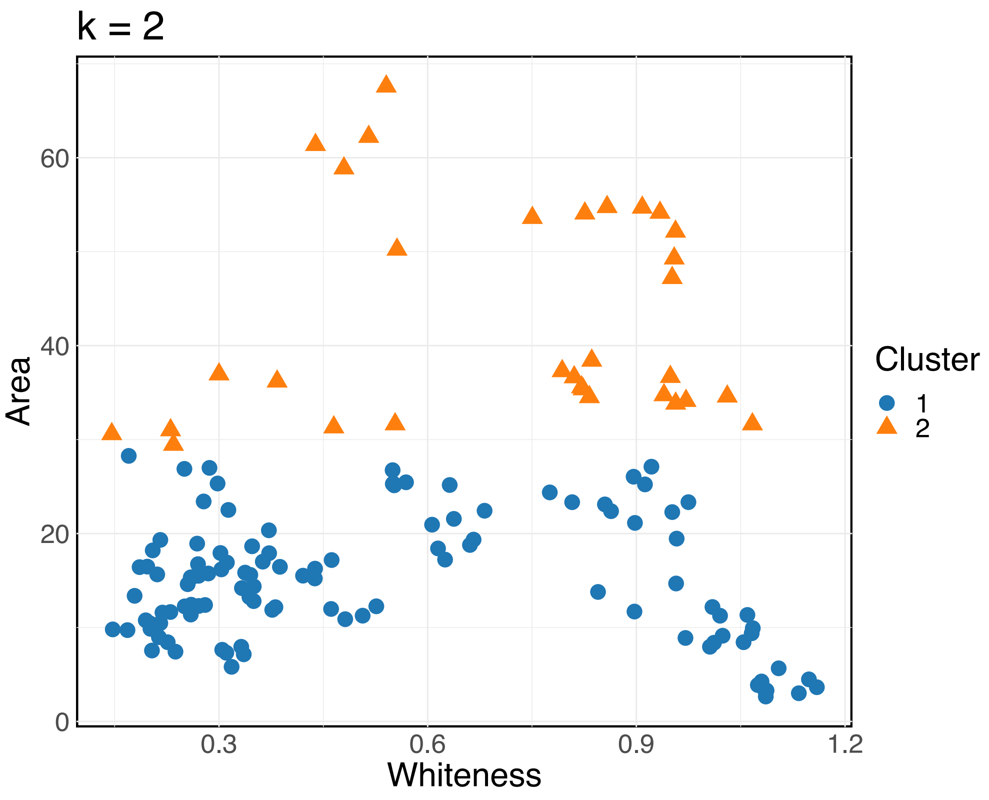
